# Interindividual variability and lateralization of µ-opioid receptors in the human brain

**DOI:** 10.1101/821223

**Authors:** Tatu Kantonen, Tomi Karjalainen, Janne Isojärvi, Pirjo Nuutila, Jouni Tuisku, Juha Rinne, Jarmo Hietala, Valtteri Kaasinen, Kari Kalliokoski, Harry Scheinin, Jussi Hirvonen, Aki Vehtari, Lauri Nummenmaa

**Affiliations:** Turku PET Centre, University of Turku, Finland; Turku PET Centre, Turku University Hospital, Finland; Department of Endocrinology, Turku University Hospital, Finland; Division of Clinical Neurosciences, Turku University Hospital, Finland; Department of Psychiatry, University of Turku and Turku University Hospital, Finland; Department of Neurology, University of Turku, Finland; Helsinki Institute for Information Technology HIIT, Department of Computer Science, Aalto University, Finland; Department of Psychology, University of Turku, Finland

## Abstract

The brain’s µ-opioid receptors (MORs) are involved in analgesia, reward and mood regulation. Several neuropsychiatric diseases have been associated with dysfunctional MOR system, and there is also considerable variation in receptor density among healthy individuals. Sex, age, body mass and smoking have been proposed to influence the MOR system, but due to small sample sizes the magnitude of their influence remains inconclusive. Here we quantified *in vivo* MOR availability in the brains of 204 individuals with no neurologic or psychiatric disorders using positron emission tomography (PET) with tracer [^11^C]carfentanil. We then used Bayesian hierarchical modeling to estimate the effects of sex, age, body mass index (BMI) and smoking on [^11^C]carfentanil binding potential. We also examined hemispheric lateralization of MOR availability. Age had regionally specific effects on MOR availability, with age-dependent increase in frontotemporal areas but decrease in amygdala, thalamus, and nucleus accumbens. The age-dependent increase was stronger in males. MOR availability was globally lowered in smokers but independent of BMI. Finally, MOR availability was higher in the right versus the left hemisphere. The presently observed variation in MOR availability may explain why some individuals are prone to develop MOR-linked pathological states, such as chronic pain or psychiatric disorders.

## Introduction

Endogenous opioids modulate multiple physiological and homeostatic functions. The most studied opioid receptors are µ-opioid receptors (MORs), which are widely expressed in the central nervous system (CNS) acting as important mediators for analgesia and reward^1^. Endogenous opioids also regulate mood^2^, social behavior^3^ and endocrine function^4, 5^. Dysregulation of the MOR system has been documented in disorders including major depression^6, 7^, schizophrenia^8^, post-traumatic stress disorder^9^, drug addiction^10^ and obesity^11^.

Opioid receptor density varies substantially between healthy humans^12, 13^, and this variation may contribute to etiology of different psychiatric conditions as well as treatment responses. First, a polymorphism in the MOR-coding gene OPRM1 influences cerebral MOR availability^14^. Second, one positron emission tomography (PET) study suggests that age and sex explain some of this variation: MOR availability increases with advancing age in cortical areas, whereas women have higher MOR availability compared to men during the reproductive years^15^. Third, lifestyle choices and health-related behaviour also affect the MOR system: Morbidly obese individuals have globally downregulated MORs^11^, and smoking has been associated with altered MOR availability^16–18^. However, interpretation of these effects is complicated by small sample sizes (typically 20–40) in neuroreceptor PET studies.

Poor replicability of neuroimaging findings has attracted attention during recent years^19^. Small sample size^20^, ubiquitous ‘researcher degrees of freedom’^21^, insufficient correction for multiple comparisons^22^, and uncertain measurements^23^ have been identified to be among the underlying reasons. Use of larger samples^19, 24^ and stricter thresholds for statistical significance^25^ have been called for to solve the problem. It has also been argued that Bayesian hierarchical modeling could facilitate reproducible research by limiting the number of paths a researcher can take in their analyses^26^ and by removing the need to correct for multiple comparisons^27^.

Here we used Bayesian hierarchical modeling with varying slopes and intercepts for different brain regions to estimate the effects of sex, age, body mass index (BMI) and smoking on cerebral MOR availability, and cerebral lateralization of the MOR system in a large sample of 204 historical subjects scanned with PET using [^11^C]carfentanil, a highly selective MOR-agonist tracer.

## Material and Methods

### Subjects

The data were retrieved from the AIVO database (http://aivo.utu.fi) of *in vivo* molecular images hosted by Turku PET Centre. We identified all the individuals with no neurologic or psychiatric disorders who had been scanned with [^11^C]carfentanil PET between 2003 and 2018 in baseline condition. The resulting data from 204 individuals consists of scans from 11 research projects and five PET scanners. No subjects abused alcohol or illicit drugs or used medications affecting the CNS. 13 females were smokers, while no males smoked. The characteristics of the subjects are summarized in Table 1. Information about the PET scanners, smoking status and handedness is summarized in Supplementary Table 1.

**Table 1.** Characteristics of the study sample.

|  | Males (n = 132) |  |  | Females (n = 72) |  |  |
| --- | --- | --- | --- | --- | --- | --- |
|  | mean | s.d. | range | mean | s.d. | range |
| Age (years) | 27.8 | 8.0 | 20–58 | 40.7 | 10.3 | 20–59 |
| Weight (kg) | 78.0 | 9.3 | 53–110 | 76.5 | 23.5 | 47–144 |
| Height (cm) | 180.7 | 5.9 | 165–198 | 165.8 | 5.7 | 151–180 |
| BMI (kg/m<sup>2</sup>) | 23.9 | 2.7 | 18–34 | 27.8 | 8.1 | 18–49 |
| Injected activity (MBq) | 297.2 | 93.5 | 218–533 | 341.1 | 109.0 | 199–519 |

### Image processing and binding quantification

Preprocessing of the PET data as well as its kinetic modeling were done using Magia^28^ (https://github.com/tkkarjal/magia), an automated processing pipeline developed at the Turku PET Centre running on MATLAB (The MathWorks, Inc., Natick, Massachusetts, United States). The preprocessing consisted of framewise realignment and coregistration of the PET and magnetic resonance images (MRIs). Tracer binding was quantified using binding potential (*BP*_*ND*_), which is the ratio of specific binding to non-displaceable binding in tissue^29^. *BP*_*ND*_ was estimated using simplified reference tissue model^30^, with occipital cortex serving as the reference region^31^. Regions of interest (ROIs) and reference regions were parcellated for each subject using FreeSurfer (http://surfer.nmr.mgh.harvard.edu/). The bilateral ROIs consisted of MOR-rich regions: amygdala, caudate, dorsal anterior cingulate cortex, inferior temporal gyrus, insula, middle temporal gyrus, nucleus accumbens, orbitofrontal cortex, pars opercularis, posterior cingulate cortex, putamen, rostral anterior cingulate cortex, superior frontal gyrus, temporal pole, and thalamus. Parametric *BP*_*ND*_ images were also calculated for full-volume analysis. They were spatially normalized to MNI-space via segmentation of T1-weighted MRIs and smoothed with an 8-mm Gaussian kernel.

### Statistical analysis

#### Model specification and estimation

Bayesian hierarchical modeling was used to estimate the effects. The models were specified using the R package BRMS^32^ that uses the efficient Markov chain Monte Carlo sampling tools of RStan (https://mc-stan.org/users/interfaces/rstan). We used weakly informative priors: For intercepts, we used the default of BRMS, i.e. Student’s t-distribution with scale 3 and 10 degrees of freedom. For predictors, a Gaussian distribution with standard deviation of 1 was used to provide weak regularization. The BRMS default prior half Student’s t-distribution with 3 degrees of freedom was used for standard deviations of group-level effects; BRMS automatically selects the scale parameter to improve convergence and sampling efficiency. The BRMS default prior LKJ(1) was used for correlations of group-level random effects.

The ROI-level models were estimated using three chains, each of which had 2000 warmup samples and 10000 post-warmup samples, thus totaling 30000 post-warmup samples. The sampling parameters were slightly modified to facilitate convergence (*adapt_delta* = 0.95; *max_treedepth* = 15). The sampling produced no divergent iterations and the Rhats were all 1.0, suggesting that the chains converged successfully. Before model estimation, continuous predictors were standardized to have zero mean and unit variance, thus making the regression coefficients comparable across the predictors. The scripts used for analyzing the data are available in https://github.com/tkkarjal/mor-variability.

#### Model comparison

We specified seven linear models with varying slopes and intercepts for the ROIs (Supplementary Figure 1). All models included subject-specific intercepts to control for variables not explicitly included in the models and dummy-covariates to control for biases between different scanners. The first model included all variables of interest (sex, age, BMI, and smoking) and sex-interactions for age and BMI; sex-smoking interaction could not be estimated because all smokers were female. The remaining models were submodels of the first model. Binding potentials were log-transformed because posterior predictive checking^33, 34^ indicated that log-transformation significantly improves model fit (see Supplementary Figure 2). The log-transformation essentially switches the model from additive to multiplicative; it also helps in model fitting because the assumption of linear additivity works poorly when the dependent variable is restricted to positive values^35^.

Predictive performance of the seven models was compared using Bayesian 10-fold cross-validation^36^, as implemented in the R package LOO (https://CRAN.R-project.org/package=loo). The subsamples used in cross-validation were created by randomly removing 10 % of the subjects. Predictive accuracy was then assessed for the leftover subjects. This procedure was repeated ten times, and the results were combined to select a model. According to the cross-validation criterion, the model without BMI (Model 4) had the best predictive accuracy, outperforming the other models significantly (Supplementary Table 2). BMI was thus not a relevant predictor of MOR availability. The posterior distribution obtained using the Model 4 (without BMI) is more closely investigated in the Results section.

#### Full-volume analyses

Traditional voxel-level analysis with Bayesian hierarchical modeling was inappropriate for the whole-brain analysis: With typically used voxel sizes (isotropic 1–6 mm) the number of voxels was so large that model estimation was computationally prohibitive. In turn, when downsampling the data to resolution whose analysis was computationally feasible (isotropic 10 mm), the resulting voxels extended over functionally heterogeneous tissue causing partial volume effects. We thus developed a new method for receptor-density-based subdivision of anatomical regions. The method clusters atlas-derived anatomical ROIs into smaller volumetric units based on the spatial receptor-density distribution within each ROI. Here the anatomical ROIs were defined using the AAL template, and the clustering was based on a high-resolution population [^11^C]carfentanil *BP*_*ND*_ map (i.e. average of all parametric *BP*_*ND*_ images analyzed in the study; see Figure 1). Hierarchical clustering, as implemented in MATLAB, was used (see https://github.com/tkkarjal/mor-variability for the code). The procedure makes the receptor availability of the clusters homogenous within clearly defined anatomical boundaries while keeping the number of volumetric units within computationally feasible limits (here 320). Once the clusters were defined, we calculated cluster-specific binding potentials for each subject, and fitted the Model 4 to the data, thus essentially estimating the effects in the whole brain.

**Figure 1.**
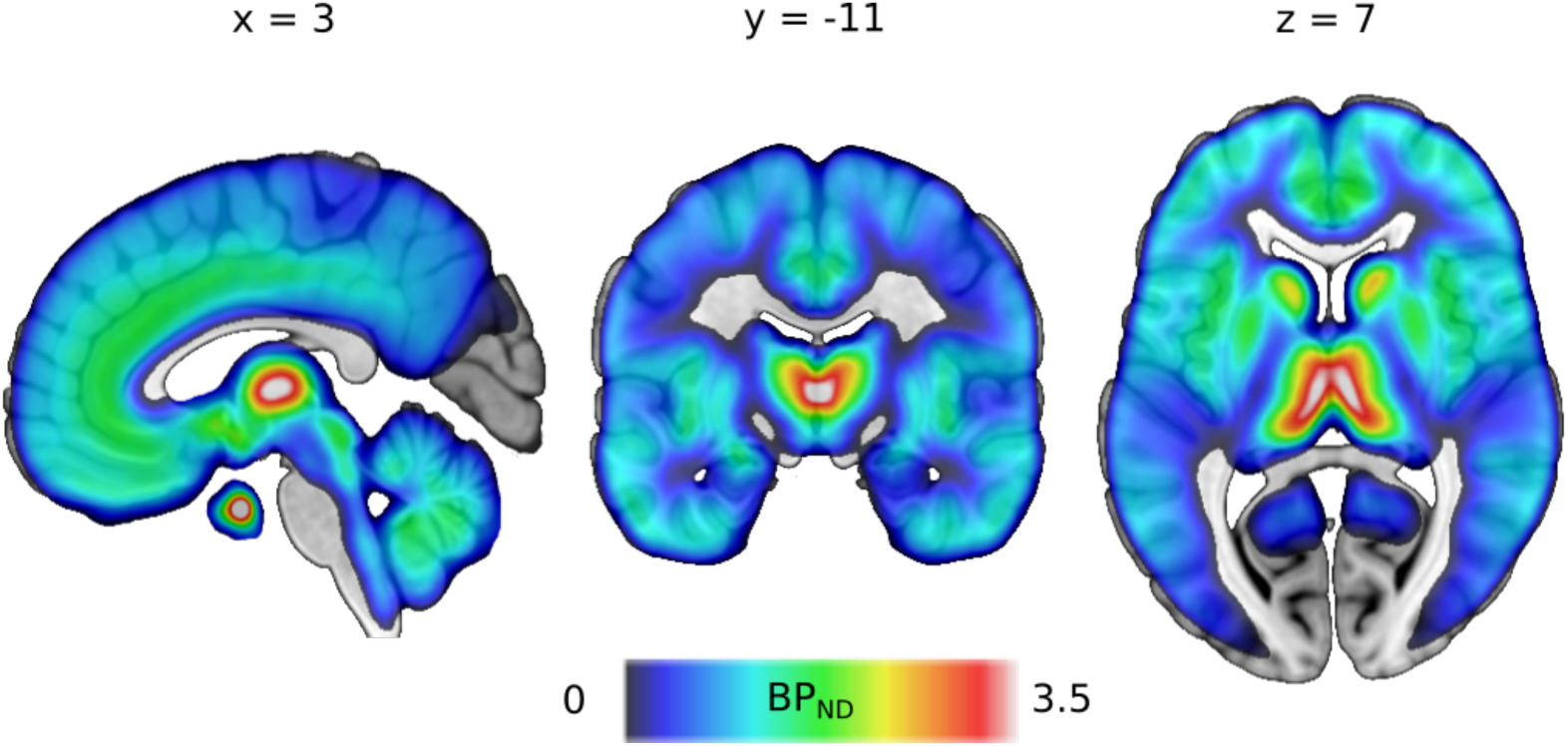
Mean distribution of µ-opioid receptors in the human brain based on the 204 [^11^C]carfentanil *BP*_ND_ images.

#### Hemispheric asymmetry

Hemispheric asymmetry was analyzed by comparing within-subject differences in binding potentials between the hemispheres. We first calculated the difference between the ROI-specific right and left hemisphere estimates. We then modeled these differences using ROI-specific intercepts using the default priors of BRMS. The effects were calculated separately for males and females.

## Results

In both sexes, MORs were widely expressed in the brain (Figure 1), consistent with previous studies^37^. Unthresholded atlases of average receptor densities for males, females and the whole sample are available at https://neurovault.org/collections/GCELSAIA/.

### Effects of age and sex

We observed regionally varying effects of age on MOR availability across both sexes (Figure 2). MOR availability decreased with ageing in amygdala, thalamus, nucleus accumbens, and cerebellum, whereas it increased with ageing in temporal regions and frontal cortex. The positive associations were stronger in males. In males, MOR availability increased with age also subcortically in putamen and insula. The proportional changes of *BP*_*ND*_ as a result of ageing 10.8 years (one standard deviation of the sample’s age distribution) on MOR availability in the ROIs are presented in Supplementary Table 3. Age-dependent sex-differences in MOR availability are visualized in Figure 3. In almost all ROIs, the mean MOR availability was higher in 20-year-old females compared to 20-year-old males (Supplementary Table 3). Because MOR availability increased with age in males faster than in females, the sex-differences decreased until around 30 years of age, after which MOR availability in males increased above females in most brain regions. Amygdala, thalamus, nucleus accumbens, and temporal pole displayed no notable sex differences at any age.

**Figure 2.**
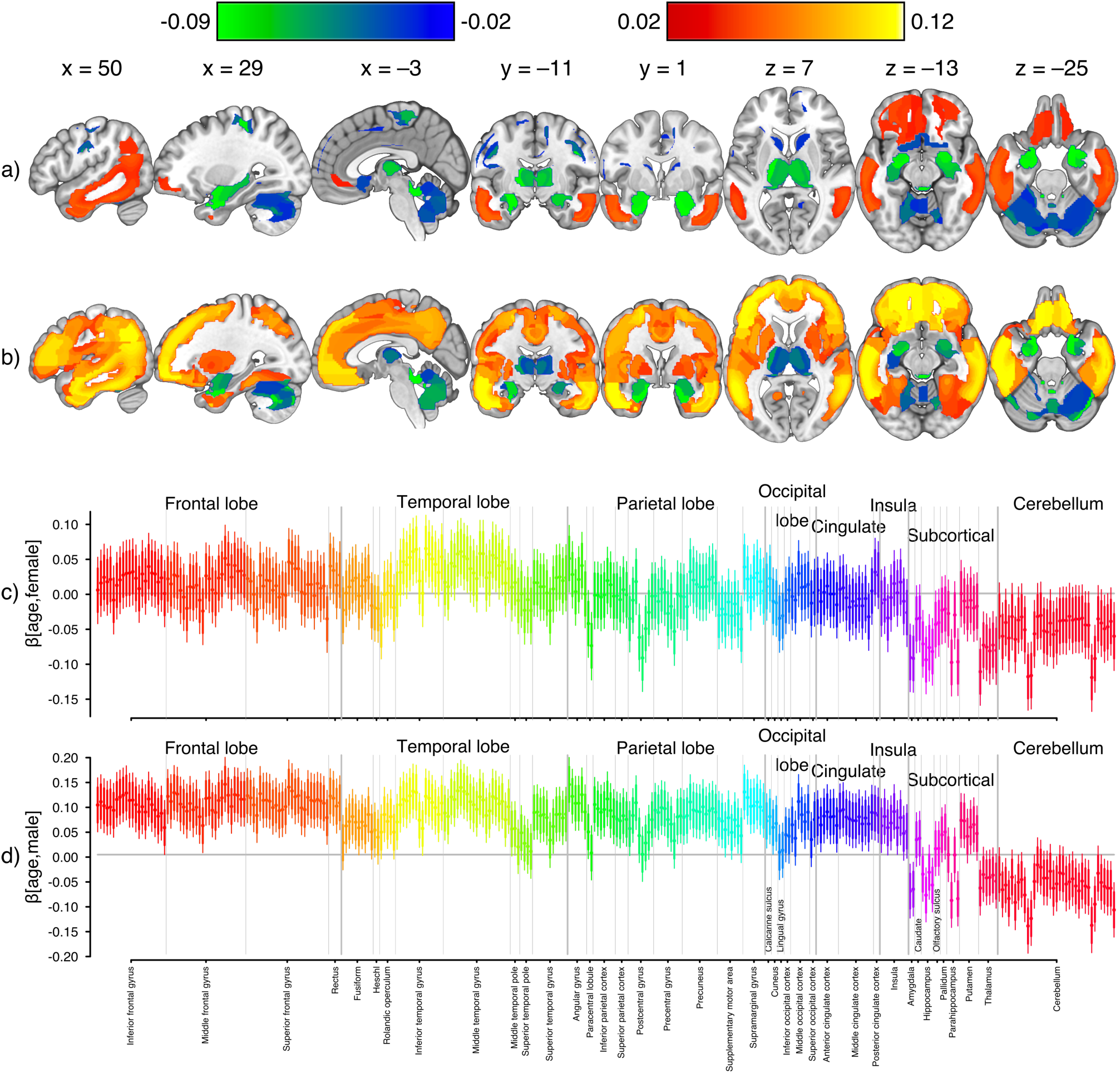
Effects of age and sex on µ-opioid receptor (MOR) availability in the brain. (a-b) Brain regions where MOR availability is associated with age in females (a) and males (b). Shown clusters have 80 % posterior interval excluding zero and absolute value of the regression coefficient at least 0.02. Colouring indicates increase (hot colours) and decrease (cool colours) of MOR availability as a function of ageing. Unthresholded maps are available at https://neurovault.org/collections/GCELSAIA/. (c-d) Summary of the posterior distributions of all clusters for the regression coefficient of age for females (c) and males (d). The filled circles represent posterior means, the thick lines 80 % posterior intervals, and the thin lines 95 % posterior intervals. Colour coding is used for highlighting the anatomical grouping of the clusters (see labels on x-axis).

**Figure 3.**
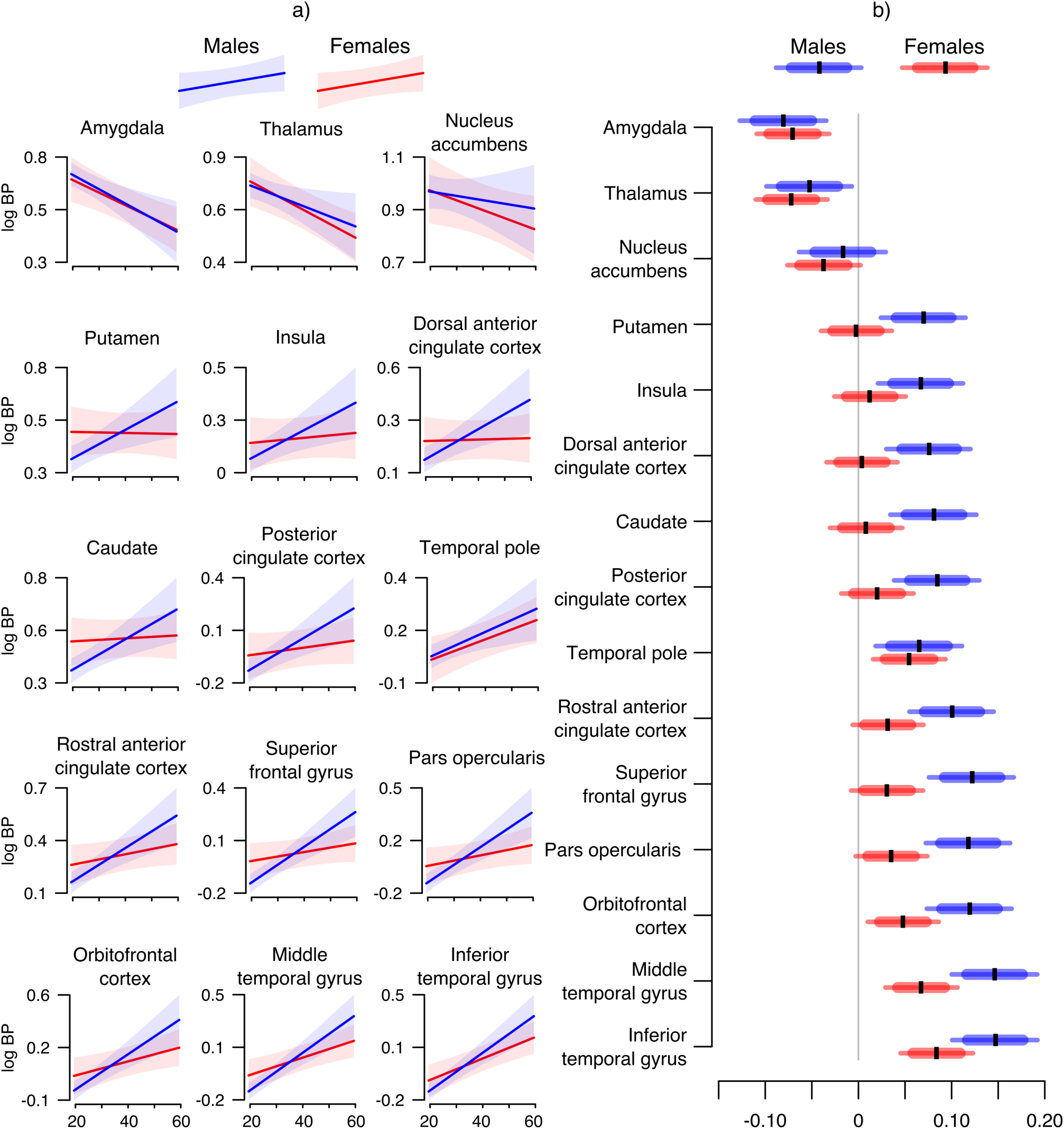
Sex-dependent ageing effects. (a) Age-dependent mean log binding potential along with 95 % highest density posterior intervals calculated over the range of data points for all regions of interest separately for males (blue) and females (red). (b) Posterior intervals of the regression coefficients of age for males (blue) and females (red). The black vertical lines represent posterior means, the thick horizontal bars 80 % posterior intervals, and the thin horizontal bars 95 % posterior intervals. The data are sorted by posterior means.

We also explored nonlinear effects of age on MOR availability in the ROIs (Supplementary Figure 3). These analyses confirmed that MOR availability decreases linearly in amygdala and thalamus, and the increase in MOR availability is steeper in males compared to females in most brain regions. In males, however, the increase was linear only until approximately 30 years of age, after which it slowed down and saturated by 50 years of age. In temporal and frontal cortices, also females exhibited similar nonlinear age-effects.

### Effects of smoking

Smokers had globally lowered binding compared to nonsmokers (Supplementary Figure 4). The reductions, ranging from 8 % to 14 % (Supplementary Table 3), were most prominent in subcortical regions such as amygdala and striatum. Because all the smokers were female, we also estimated the effects of smoking using female-only data. This analysis yielded similar results.

### Hemispheric asymmetry

The posterior distributions for absolute differences between regional binding potentials in the right and left hemispheres are presented in Figure 4. All differences were within +/− 6 % (Supplementary Table 4). In most brain regions, including thalamus, caudate, cingulate cortex, orbitofrontal cortex, and putamen, MOR availability was higher in the right hemisphere. Only in nucleus accumbens and amygdala was MOR availability higher in the left hemisphere. This lateralization was consistent across the sexes. We also examined hemispheric differences in subsamples of the 128 right-handed and the seven left-handed individuals, and found that both groups display lateralization towards the right hemisphere.

**Figure 4.**
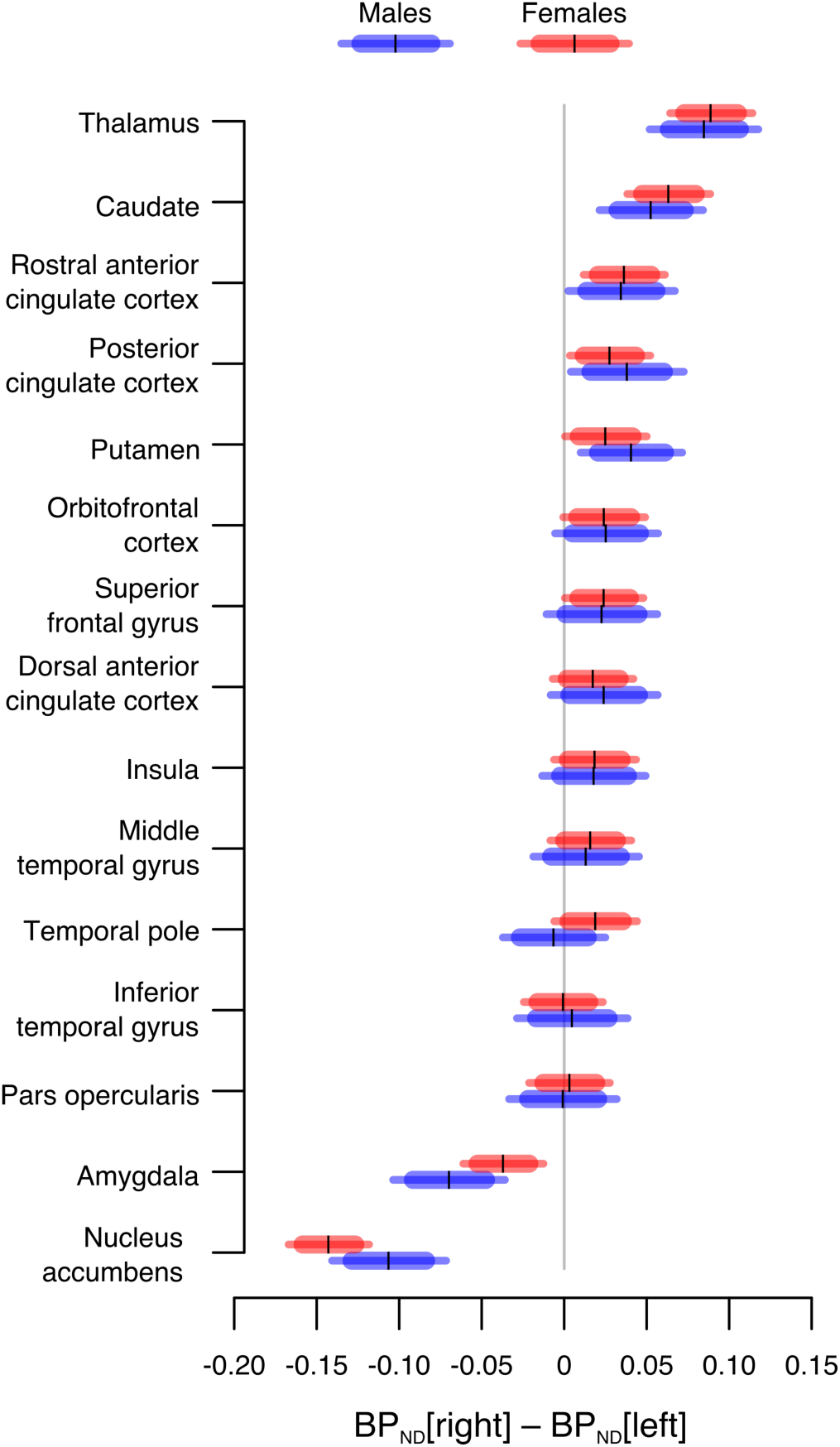
Posterior distribution of hemispheric differences estimated in each region separately for females (red) and males (blue). Distributions located right to the dashed line have higher binding in the right hemisphere and vice versa. The black vertical lines represent posterior means, the thick horizontal bars 80 % posterior intervals, and the thin horizontal bars 95 % posterior intervals.

## Discussion

Our main findings were that sex, age, and smoking influence [^11^C]carfentanil binding potential in the brain. BMI was not associated with MOR availability. In most brain regions, MOR availability was slightly higher in the right than in the left hemisphere.

### Effects of age on endogenous opioid system

Effects of ageing on MORs were regionally dependent. In both sexes, advancing age was associated with reductions in [^11^C]carfentanil *BP*_*ND*_ in thalamus, amygdala, nucleus accumbens, and cerebellum. On the contrary, MOR availability in frontotemporal areas increased with age in both sexes. Nonlinear models revealed that this upregulation reaches a plateau at around 40–50 years of age. The age-related increase in cortical MOR availability is consistent with previous data from autoradiography studies^12, 13, 38^. Also, the only prior PET study that has investigated effects of ageing on MOR availability in a significantly smaller sample reported positive association in cortical areas^15^. Our data thus support cortical increase of MOR availability with advancing age, but highlight that the increase saturates at some point and that the effect is reversed in some brain structures.

The age-related decrease in MOR availability observed in thalamus, amygdala and nucleus accumbens is consistent with the general pattern of ageing-related brain atrophy, emerging as reduced cortical thickness and reduced grey and white matter volume^39^. Many other receptor systems, including serotonergic and dopaminergic systems, undergo receptor loss with advancing age^40, 41^. The decrease in MOR availability likely reflects similar receptor loss. This may have implications for chronic pain. In PET studies, fibromyalgia and central neuropathic pain are associated with reduced opioid receptor availability in nucleus accumbens, amygdala and thalamus^42, 43^. These brain regions also display enhanced opioidergic processing during acute pain^44^. Age-related reduction in MOR density in these pain modulatory brain regions could thus reflect lowered pain-attenuation capacity, possibly explaining some of the increased prevalence of pain disorders among the elderly^45^.

While consistent with prior findings, the age-related *increase* in cortical MOR availability is in contrast with the age-related neural atrophy. The increase in MOR availability likely reflects increase in receptor density, as data from rodents indicates that MOR density increases with age^46^, and human autoradiography studies suggest that the age-related increase in cortical MOR availability results from increased receptor density rather than affinity^12, 13^. As ageing also results in decreased endogenous opioid concentration in rodent CNS^47^, it is possible that the receptors are upregulated to compensate for the decreased endogenous opioid drive^48^.

### Sex differences in µ-opioid receptor availability

Regional effects of ageing on MOR were stronger in males, and also observed more widely in the brain than in females. Previously, two PET studies have investigated sex differences in cerebral MOR availability. Data from the first study suggest that during their reproductive years females have higher MOR availability than males in thalamus, caudate, and amygdala, but that the difference disappears or reverses after menopause^15^. In the second study with age-matched 40-year-old participants, males had higher MOR availability in cingulate cortex and ventral striatum^49^. The present large-scale dataset suggests that the sex difference in MOR availability is actually *reversed* during ageing: In the reproductive age women have higher MOR availability than men in multiple brain regions, whereas older males have higher MOR availability in cortical and most subcortical areas.

These sex differences might be linked with gonadal steroids: In rodents, subcutaneous injection of estrogen increases MOR mRNA levels in the CNS^50^. Human studies have shown that in females the estrogen-state influences MOR availability in thalamus, amygdala and nucleus accumbens^51^. Menopause and the associated low-estrogen state in ageing females might thus explain some of the observed sex differences. Also in males, the estrogen-to-testosterone ratio increases with age^52^, and the elevated estrogen tone might in turn upregulate the MOR system.

In some brain regions, the age-dependencies were nonlinear. In males, MOR availability increased rapidly between the ages 20 and 30, after which the increase slowed down and plateaued at around 40–50 years. Similar saturation pattern was observed in females in frontal and temporal cortices. These data indicate that the linear sex-interactions that were observed in temporal cortex could result from differences in the age distributions rather than from biological sex differences. Elsewhere in the brain, the steeper increase of MOR availability in males is more likely to be of biological origin.

### Smoking and endogenous opioid system

We found that smokers had globally lowered MOR availability. The effect was particularly large in striatum, insula and amygdala. MOR availability reductions in smokers have been previously documented in many brain areas, including anterior cingulate cortex, thalamus, amygdala, insula and striatum^16, 17^. Our data confirm these reductions. Nicotine administration induces release of endogenous opioids^53^, a mechanism that presumably underlies nicotine-induced reward^54^. Regular smoking thus frequently activates the endogenous opioid system, and our data suggest that it leads to downregulation of MORs, possibly to prevent overactivation of the MOR system.

### Body mass index in the non-obese range does not correlate with µ-opioid receptor availability

BMI was not associated with [^11^C]carfentanil *BP*_*ND*_. This was against our expectations, since MOR availability is decreased in morbidly obese subjects^11, 55^. Furthermore, binge eating disorder (BED) patients also have reduced MOR availability in several brain regions^56^. Because our subjects were mostly normal-weight, BMI may be associated with MOR only in the pathological (obese) end of the spectrum. BMI is widely used as a marker for obesity and excess fat, but it does not directly measure body composition or obesity, nor does it necessarily reflect the changes of body adiposity arising from factors such as subject’s age or muscle mass^57^. Lowered MOR availability in the morbidly obese and BED patients may therefore result from pathological eating or dysfunction of the pleasure system rather than from high body mass itself.

### Lateralization of endogenous opioid system

We found differences of 1–5 % between the binding potentials of the two hemispheres. *BP*_*ND*_ was higher in the right than in the left hemisphere in most brain areas. Only in nucleus accumbens and amygdala *BP*_*ND*_ was lateralized to the left hemisphere. Handedness did not explain the direction of lateralization. Enkephalin levels have been observed to differ in post-mortem tissue samples from anterior cingulate cortex^58^, showing that opioid peptide concentrations may differ between the hemispheres, possibly explaining some of the lateralization.

Previous studies have revealed functional asymmetry in amygdala’s emotion processing. Under direct electrical stimulation, right amygdala produces only negative emotions, whereas left amygdala induces both positive and negative emotions^59^. In PET studies, depression has been connected to lowered MOR availability in amygdala^6^, and acute positive mood is associated with amygdala’s MOR system activation^60^. Amygdala has important excitatory effects on ipsilateral nucleus accumbens, facilitating motivated behaviour^61^. Nucleus accumbens and amygdala were the only regions where the MOR system was lateralized to the left hemisphere. MOR lateralization in amygdala and nucleus accumbens might reflect lateralization of emotion processing, the left amygdala with higher MOR availability being more responsible for emotional processes.

Thalamus showed the most prominent rightward lateralization with 5 %. Thalamus relays somatosensory and nociceptive signals to cerebral cortex^62^. Acute pain is associated with endogenous opioid release in thalamus, presumably to alleviate pain^44^. Right thalamus may be more specialized to pain processing than the left, since following a stroke and brain damage, thalamic pain syndrome develops more frequently if the thalamic lesion has been right-sided^63^. Our finding of increased MOR availability in the right thalamus is consistent with lateralized central pain processing.

### Limitations

The primary limitation of the study is that the data were sampled from 11 distinct study projects and obtained using five different PET scanners. Even if the scanners are cross-calibrated, they may produce slightly different *BP*_*ND*_ estimates. We however corrected for potential scanner-related biases in the analyses. The available data was not ideally detailed – in females, age of the possible menopause or the phase of menstrual cycle was not recorded. Our sample had subjects with 20–60 years of age, and the conclusions may not be generalizable outside this range. We had no information about the genetic profile of our subjects, and we could thus not assess the influence of genetic factors. Furthermore, the data set was not fully representative for all subgroups of interest. For example, there were only seven confirmed left-handed subjects and 13 smokers, and all smokers were females. Another limitation is related to the concept of binding potential: *BP*_*ND*_ is the product of i) density of the target receptors unoccupied by endogenous ligands and ii) affinity of the radioligand to the receptor^1, 64^, and in a single PET scan it is impossible to differentiate these components. [^11^C]carfentanil is an agonist tracer preferably binding in MORs in the high-affinity state^1^. While endogenous opioids compete for binding sites with [^11^C]carfentanil^65 66^, at least in rats the basal opioid concentrations are low^67^. Accordingly, [^11^C]carfentanil *BP*_*ND*_ in baseline condition likely reflects mostly density of high-affinity MORs.

## Conclusions

Age, sex, and smoking influence cerebral MOR availability. Ageing was associated with decreased MOR availability in thalamus and amygdala, possibly reflecting age-associated neurodegeneration and pain burden. In temporal and frontal regions, MOR availability increased with ageing, which might reflect a compensatory process to reduced endogenous opioid concentrations. MOR availability increased with age faster in males than in females, and these sex differences may pertain to sex-specific hormonal modulation of MOR system. Smoking was associated with globally decreased MOR availability, likely resulting from downregulation of MORs as a consequence of continuous activation of the MOR system. Even if alterations in MOR system have been reported in obesity and binge eating, BMI was not associated with MOR availability. We found that the human MOR system manifests consistent but minor asymmetry between the two hemispheres. From experimental point of view, these data highlight that studies with [^11^C]carfentanil need to have well-matched, non-smoking samples and single-sex subject selection, unless sufficient statistical power for analysing sex differences can be achieved. Given the prominent effect of ageing on MORs especially before the age of 30, even small age differences between tested groups can lead to artificial group differences. Altogether, these data show that healthy humans vary significantly in their cerebral MOR availability, and that age, sex and smoking status explain a part of this variation. Variability in MOR system might be one neurobiological mechanism explaining why some individuals are more vulnerable to develop MOR-linked pathological states such as chronic pain or neuropsychiatric disorders.

## Acknowledgements

This study was supported by Academy of Finland (grant #294897# to LN) and Sigrid Juselius Foundation. We thank the Finnish Cultural Foundation (Southwest Finland Fund) for personal grants to T. Kantonen and T. Karjalainen, as well as State Funding for University-level Health Research given to T. Karjalainen. We also thank the Emil Aaltonen Foundation for personal grant to T. Kantonen. This article has been posted on preprint server bioRxiv (https://www.biorxiv.org).

