## Supplementary material for "Interindividual variability and lateralization of µ-opioid receptors in the human brain"

i. PET scanners, smoking status and handedness

ii. Compared models

iii. Posterior predictive distributions with and without log-transformation

iv. Model comparison results

v. Effect size estimates for sex differences, ageing, and smoking

vi. Nonlinear effects of age

vii. Effects of smoking on MOR availability

viii. Relative differences between the hemispheres

**PET scanners, smoking status and handedness**

Supplementary Table 1. The information about PET scanners, smoking status and handedness of the sample.

| | Males (n = 132) | Females (n = 72) |
|---|---|---|
| PET scanner* (n) | | |
| GE Advance | 6 | 0 |
| HRRT | 31 | 34 |
| PET/CT | 30 | 0 |
| GE Discovery | 10 | 38 |
| PET/MR | 55 | 0 |
| Smoking status** (n) | | |
| Smoking | 0 | 13 |
| Nonsmoking | 129 | 53 |
| Unknown | 3 | 6 |
| Handedness (n) | | |
| Right | 80 | 48 |
| Left | 5 | 2 |
| Ambidextrous | 0 | 1 |
| Unknown | 47 | 21 |

*PET scanner abbreviations: GE Advance (GE Advance, GE Healthcare), HRRT (HRRT, Siemens Medical Solutions), PET/CT (GE Discovery VCT PET/CT, GE Healthcare), GE Discovery (Discovery 690 PET/CT, GE Healthcare), PET/MR (Ingenuity TF PET/MR, Philips).

**Nine individuals whose smoking status was unknown were classified as nonsmokers because it enabled us to estimate all effects using a single model, and as in Finland, less than 20 % of the adult population smoked cigarettes in 2000 (http://urn.fi/URN:NBN:fi-fe2018102938947).

**Compared models**

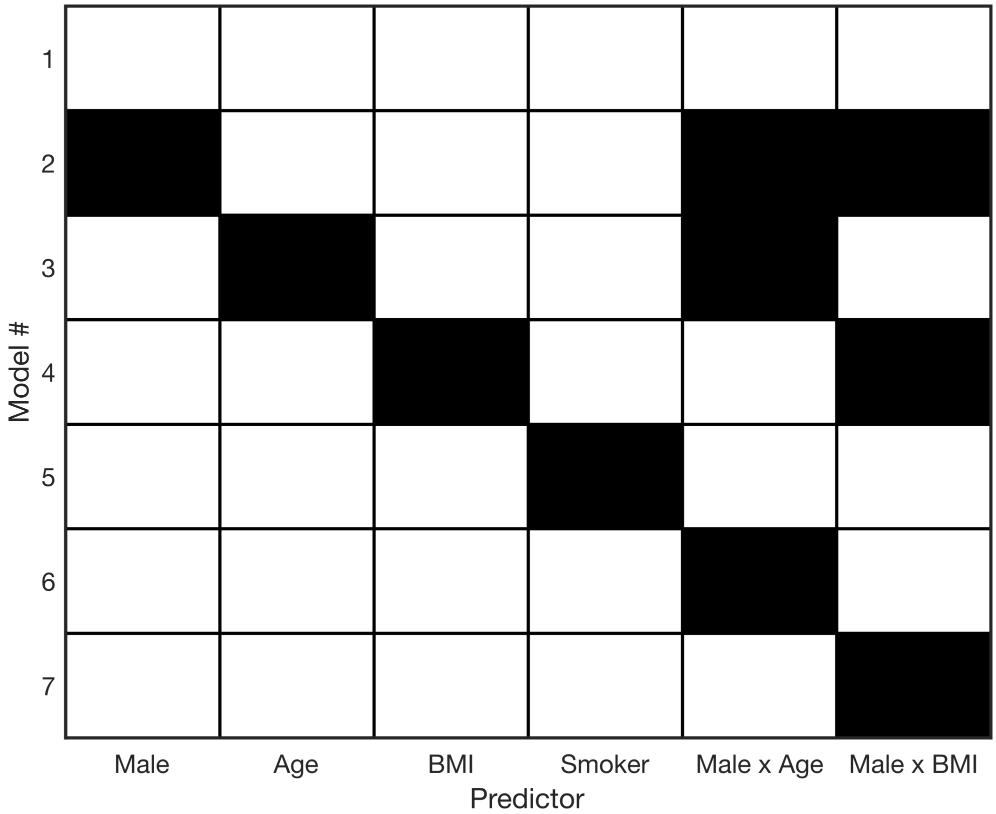


Supplementary Figure 1. The specified models. Columns represent variables included in the models. Each row thus shows white variables which were included in the model. Black squares represent omitted variables.

**Posterior predictive distributions with and without log-transformation**

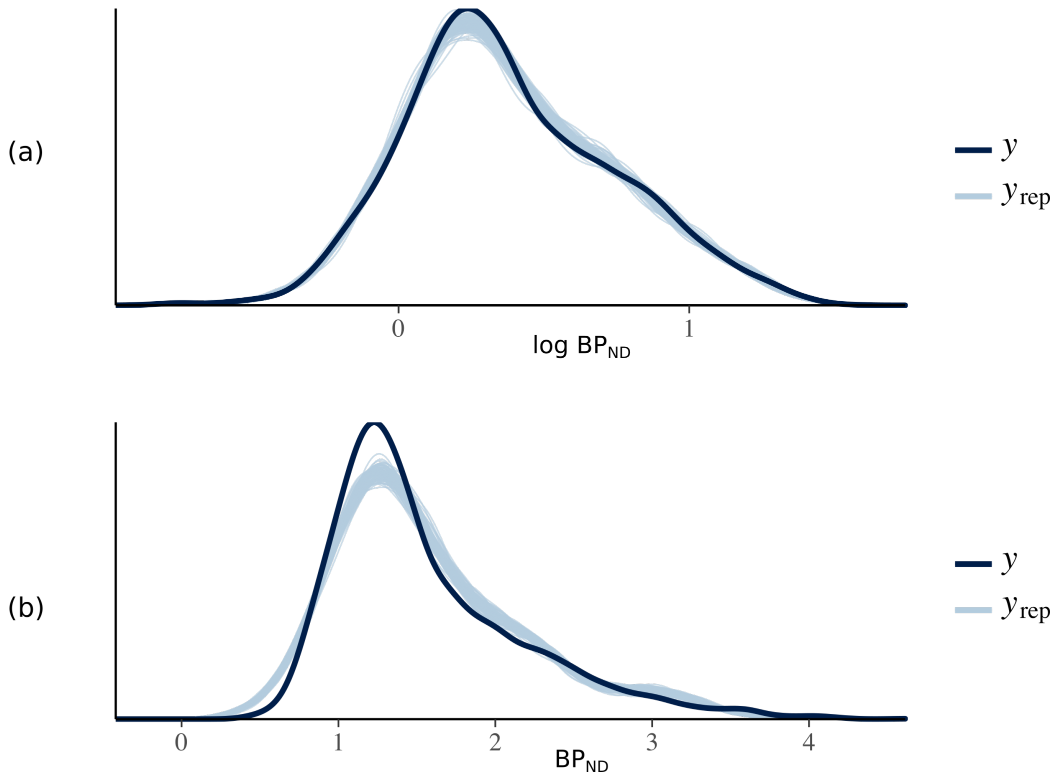


Supplementary Figure 2. Kernel density of the observed (black) and estimated (blue) data. Shown are 100 draws. Log-transformation, shown in top row (a), of binding potential resulted in significantly better fit compared to non-transformed binding potential, shown in (b).

### Model comparison

The model comparison results are summarized in Supplementary Table 2. According to Bayesian 10-fold cross-validation, the Model 4 (without BMI) had the highest predictive accuracy. In other words, all the other predictors except BMI significantly improved predictive accuracy of regional $BP_{ND}$. Accordingly, posterior distribution of Model 4 was investigated in detail.

Supplementary Table 2. Model comparison results. The table shows the expected predictive accuracies of the models, as quantified by expected log predictive density (elpd). Standard errors of the estimates are given in parentheses. The table also shows the pairwise differences in elpd. For each column, negative values mean that the corresponding model has higher predictive accuracy. For example, the last row of the column corresponding to Model 1 states that Model 7 outperforms Model 1, and the difference (7.1) is over 2 times the standard error (2.6) of the difference.

| | elpd_kfold | Difference in expected log predictive accuracy | | | | | | |
|---|---|---|---|---|---|---|---|---|
| Model number | | 1 | 2 | 3 | 4 | 5 | 6 | 7 |
| 1 | 1276.2 (52.0) | - | 11.6 (8.3) | 153.3 (16.1) | -18.5 (3.6) | 23.9 (8.9) | 32.0 (7.0) | -7.1 (2.6) |
| 2 | 1264.6 (51.5) | -11.6 (8.3) | - | 141.7 (15.2) | -30.2 (9.1) | 12.3 (13.7) | 20.4 (4.5) | -18.7 (8.3) |
| 3 | 1123.0 (50.8) | -153.3 (16.1) | -141.7 (15.2) | - | -171.8 (17.4) | -129.4 (18.1) | -121.3 (14.3) | -160.3 (16.8) |
| 4 | 1294.8 (51.9) | 18.5 (3.6) | 30.2 (9.1) | 171.8 (17.4) | - | 42.4 (9.3) | 50.5 (8.9) | 11.5 (2.7) |
| 5 | 1252.3 (52.2) | -23.9 (8.9) | -12.3 (13.7) | 129.4 (18.1) | -42.4 (9.3) | - | 8.1 (12.5) | -31.0 (9.6) |
| 6 | 1244.2 (51.7) | -32.0 (7.0) | -20.4 (4.5) | 121.3 (14.3) | -50.5 (8.9) | -8.1 (12.5) | - | -39.0 (8.0) |
| 7 | 1283.3 (51.9) | 7.1 (2.6) | 18.7 (8.3) | 160.3 (16.8) | -11.5 (2.7) | 31.0 (9.6) | 39.0 (8.0) | - |

### Effect size estimates for sex differences, ageing, and smoking

Supplementary Table 3. Effect size estimates for ageing, sex differences, and smoking, expressed in percent of change. The ageing effects represent how much the binding potential changes as a result of ageing 10.8 years, that is one standard deviation of the sample's age distribution. Sex-differences are calculated as the difference between the mean binding potentials at ages 20, 40 and 60 years. F = female, M = male, m = median, l = lower bound (2.5 % percentile), u = upper bound (97.5 % percentile).

| ROI | F aging | | | M aging | | | F > M @ 20 y | | | F > M @ 40 y | | | F > M @ 60 y | | | smoking | | |
|---|---|---|---|---|---|---|---|---|---|---|---|---|---|---|---|---|---|---|
| | m | l | u | m | l | u | m | l | u | m | l | u | m | l | u | m | l | u |
| amy | -7 | -10 | -4 | -8 | -11 | -4 | -3 | -11 | 7 | -1 | -6 | 5 | 1 | -11 | 15 | -11 | -18 | -4 |
| cau | 1 | -2 | 4 | 8 | 4 | 13 | 15 | 5 | 26 | 0 | -5 | 7 | -12 | -22 | 0 | -14 | -20 | -6 |
| dacc | 0 | -3 | 4 | 8 | 4 | 12 | 9 | 0 | 19 | -5 | -10 | 1 | -16 | -26 | -6 | -9 | -15 | -1 |
| inftemp | 9 | 5 | 12 | 16 | 11 | 20 | 8 | -1 | 18 | -4 | -9 | 2 | -14 | -24 | -3 | -8 | -14 | 1 |
| ins | 1 | -2 | 4 | 7 | 3 | 11 | 7 | -2 | 17 | -3 | -9 | 2 | -13 | -23 | -1 | -11 | -18 | -4 |
| midtemp | 7 | 4 | 10 | 16 | 11 | 20 | 12 | 2 | 22 | -3 | -9 | 2 | -16 | -26 | -5 | -8 | -15 | 0 |
| nacc | -4 | -7 | 0 | -2 | -5 | 2 | 0 | -8 | 10 | -3 | -9 | 3 | -7 | -18 | 6 | -9 | -16 | -1 |
| ofc | 5 | 2 | 8 | 13 | 8 | 17 | 9 | 0 | 19 | -4 | -9 | 1 | -16 | -26 | -5 | -10 | -16 | -3 |
| parsop | 4 | 0 | 7 | 12 | 8 | 17 | 11 | 1 | 21 | -5 | -10 | 0 | -18 | -28 | -8 | -9 | -15 | -2 |
| pcc | 2 | -1 | 5 | 9 | 5 | 13 | 8 | -2 | 18 | -4 | -10 | 1 | -15 | -25 | -4 | -8 | -15 | 0 |
| put | 0 | -3 | 3 | 7 | 3 | 11 | 13 | 3 | 24 | -1 | -7 | 5 | -14 | -23 | -2 | -12 | -18 | -5 |
| racc | 3 | 0 | 7 | 11 | 6 | 15 | 10 | 0 | 20 | -3 | -8 | 2 | -15 | -25 | -4 | -11 | -17 | -4 |
| supfront | 3 | 0 | 6 | 13 | 9 | 17 | 15 | 5 | 25 | -3 | -8 | 2 | -18 | -28 | -7 | -11 | -17 | -4 |
| tempol | 6 | 2 | 9 | 7 | 3 | 11 | -2 | -10 | 8 | -4 | -9 | 2 | -5 | -17 | 8 | -11 | -17 | -4 |
| tha | -7 | -10 | -4 | -5 | -9 | -1 | 2 | -7 | 11 | -2 | -7 | -4 | -5 | -16 | 8 | -10 | -16 | -2 |

### Nonlinear effects of age

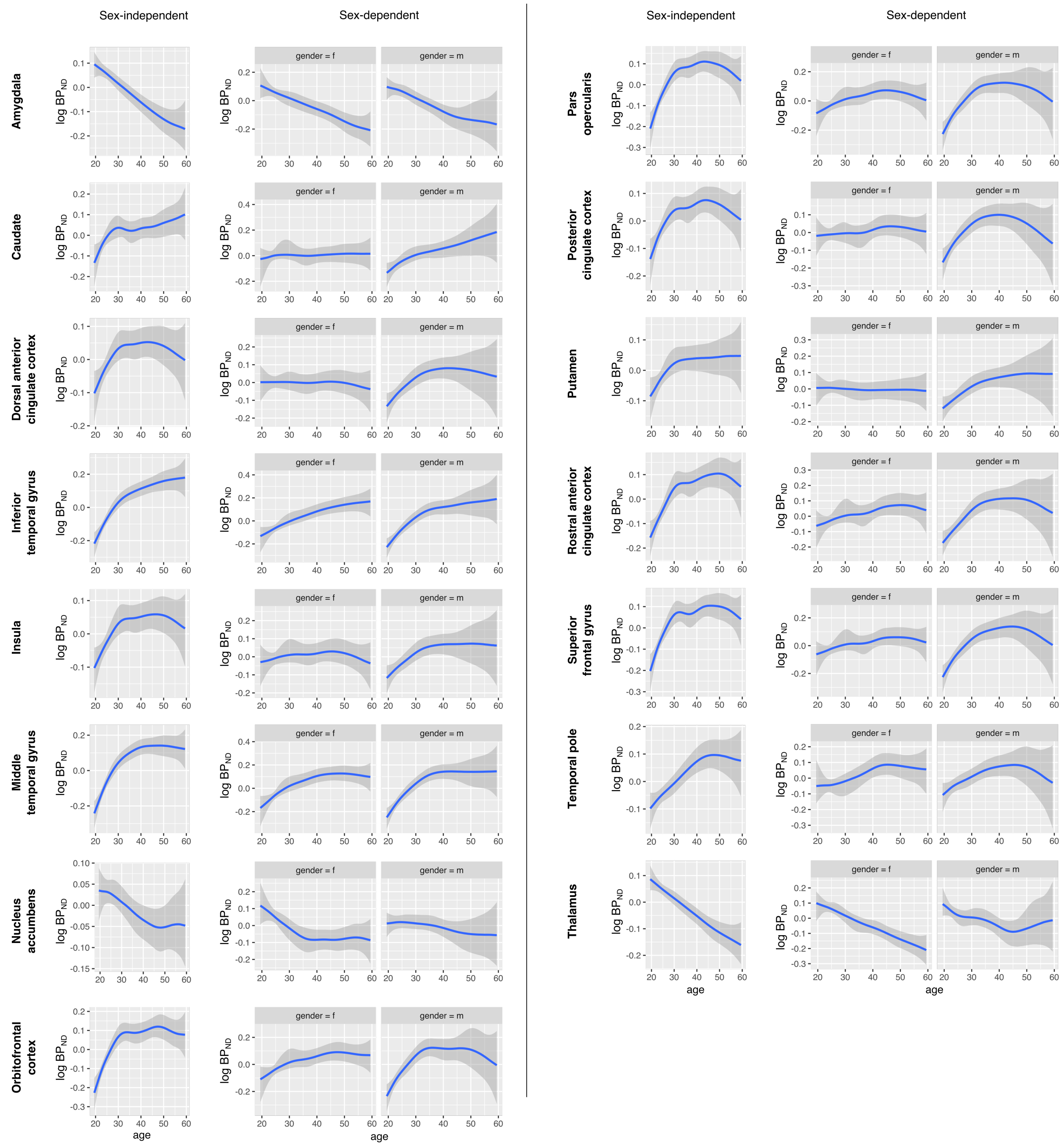


Supplementary Figure 3. Nonlinear effects of age on MOR availability in 15 ROIs. The figures show the sex-independent as well as sex-dependent nonlinear effects.

### Effects of smoking on MOR availability

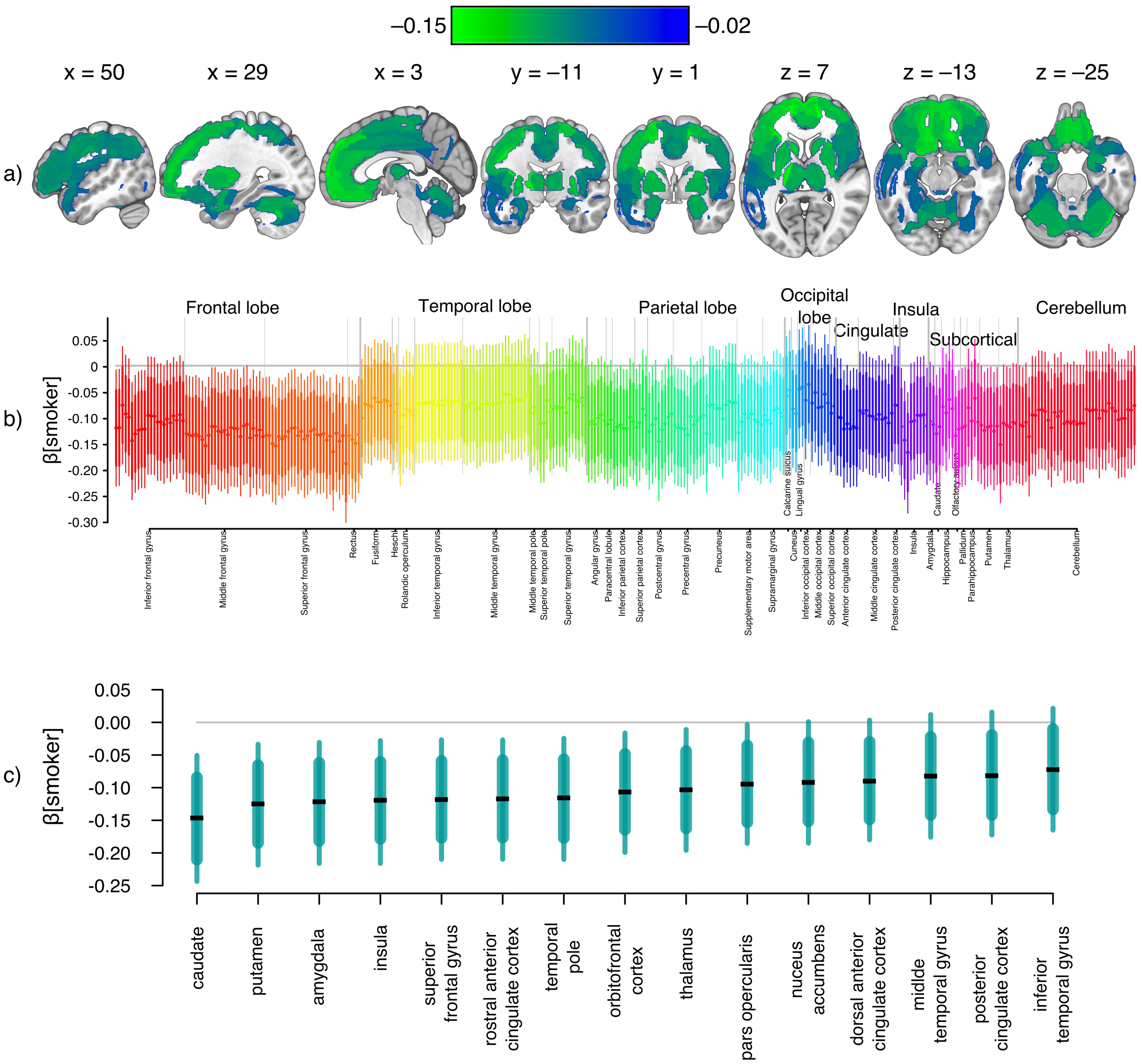


Supplementary Figure 4. Effects of smoking on MOR availability in the brain. (a) Brain regions where smokers had lowered [$^{11}$C]carfentanil $BP_{ND}$ compared to nonsmokers. Shown are clusters whose 80 % posterior interval excluded zero and in which the absolute value of the regression coefficient was at least 0.02. (b) Summary of the posterior distributions of all clusters for the regression coefficient of smoking in the whole-brain comparison. The filled circles represent posterior means, the thick lines 80 % posterior intervals, and the thin lines 95 % posterior intervals.

(c) ROI-level posterior distributions. The filled circles represent posterior means, the thick lines 80 % posterior intervals, and the thin lines 95 % posterior intervals.

**Relative differences between the hemispheres**

Supplementary Table 4. Relative difference between regional MOR availabilities in the two hemispheres. m = median, l = lower bound (2.5 % percentile), u = upper bound (97.5 % percentile).

| ROI | Females: RH - LH (%) | | | Males: RH - LH (%) | | |
|---|---|---|---|---|---|---|
| | m | l | u | m | l | u |
| amygdala | -4 | -5 | -2 | -2 | -3 | -1 |
| caudate | 3 | 2 | 5 | 4 | 3 | 6 |
| dorsal anterior cingulate cortex | 2 | 0 | 4 | 1 | 0 | 3 |
| inferior temporal gyrus | 1 | -3 | 4 | 0 | -2 | 2 |
| insula | 2 | -1 | 4 | 2 | 0 | 3 |
| middle temporal gyrus | 1 | -1 | 4 | 2 | -1 | 4 |
| nucleus accumbens | -4 | -5 | -3 | -5 | -6 | -4 |
| orbitofrontal cortex | 2 | 0 | 5 | 2 | 0 | 4 |
| pars opercularis | 0 | -3 | 3 | 0 | -2 | 2 |
| posterior cingulate cortex | 4 | 1 | 6 | 3 | 1 | 4 |
| putamen | 3 | 1 | 5 | 2 | 0 | 3 |
| rostral anterior cingulate cortex | 3 | 1 | 5 | 3 | 1 | 4 |
| superior frontal gyrus | 2 | -1 | 5 | 2 | 0 | 4 |
| temporal pole | -1 | -3 | 2 | 2 | 0 | 4 |
| thalamus | 5 | 3 | 6 | 5 | 4 | 6 |
